# Calcium binding and permeation in TRPV channels: insights from molecular dynamics simulations

**DOI:** 10.1101/2022.09.07.506889

**Authors:** Chunhong Liu, Chen Song

## Abstract

Some calcium channels selectively permeate Ca^2+^, despite the high concentration of monovalent ions in the surrounding environment, which is essential for many physiological processes. Without atomistic and dynamical ion permeation details, the underlying mechanism of Ca^2+^ selectivity has long been an intensively studied, yet controversial, topic. This study takes advantage of the homologous Ca^2+^-selective TRPV6 and non-selective TRPV1 and utilizes the recently solved open-state structures and a newly developed multi-site calcium model to investigate the ion binding and permeation features in TRPV channels by molecular dynamics simulations. Our results revealed that the open-state TRPV6 and TRPV1 show distinct ion-binding patterns in the selectivity filter, which lead to different ion permeation features. Two Ca^2+^ ions simultaneously bind to the selectivity filter of TRPV6 compared with only one Ca^2+^ in case of TRPV1. Multiple Ca^2+^ binding at the selectivity filter of TRPV6 permeated in a concerted manner, which could efficiently block the permeation of Na^+^. Cations of various valences differentiate between the binding sites at the entrance of the selectivity filter in TRPV6. Ca^2+^ preferentially binds to the central site with a higher probability of permeation, repelling Na^+^ to a peripheral site. Therefore, we believe that ion binding competition at the selectivity filter of calcium channels, including the binding strength and number of binding sites, determines Ca^2+^ selectivity under physiological conditions. Additionally, our results showed that pore helix flexibility and the cytosolic domain of TRPV channels regulate ion permeability.

## Introduction

Ion channels are important regulators of intracellular and extracellular ion concentration. The opening of ion channels generates ionic currents, which are fundamental signals in the nervous systems of animals (Hodgkin and Huxley, 1952). The selectivity of ion channels enables them to allow specific types of ions to pass through, which is a key feature of their biological function. For example, selective voltage-gated sodium channels (Na_V_ channels) and voltage-gated potassium channels (K_V_ channels) in neuronal membranes are involved in action-potential generation and conduction. These channels respond to changes in membrane potential and generate currents specifically conducted by Na^+^ or K^+^. If these channels lose their selectivity, inward sodium and outward potassium currents cannot be generated, and action potentials cannot spread among neurons (Abdul Kadir et al., 2018). Divalent ion channels, such as calcium channels that are weakly to highly selective for Ca^2+^, are also involved in essential physiological processes, including muscle contraction, neurotransmitter release, and hormone secretion (Ebashi and Endo, 1968; Ghosh and Greenberg, 1995; Wiederkehr et al., 2011). Some calcium channels are highly selective. For example, Ca_V_ channels can reach a selectivity of P_Ca_: P_Na_ ~ 1000 despite the high concentration of sodium in the extracellular environment. How ion channels distinguish between ions with subtle differences and selectively permeate specific types of ions are fundamental scientific questions in the field of ion channel research.

For ion channels with available structures, molecular dynamics (MD) simulation is a powerful method for studying the microscopic mechanism of ion permeation and selectivity. There has been much success using this method to elucidate the permeation and selectivity mechanism of monovalent cation channels, such as potassium and sodium channels (Bernèche and Roux, 2001; Jensen et al., 2010; Corry and Thomas, 2012; Xia et al., 2013; Köpfer et al., 2014; Flood et al., 2018; Kopec et al., 2018). Dehydration of the first solvation shell of cations and interactions between multiple ions are thought to be pivotal in the selectivity mechanism of these monovalent ion channels (Mironenko et al., 2021). For calcium channels, multiple hypotheses have been proposed to explain their selectivity (Boda et al., 2000; Corry et al., 2001; Sather and McCleskey, 2003). In the early 1980s, Tsien proposed that calcium channels achieve selectivity by forming multiple binding sites that have a higher affinity for Ca^2+^ than other ions in the pore. Ca^2+^ has been proposed to permeate in a knock-off manner, which requires a repulsive force from a second ion to facilitate the detachment of Ca^2+^ from the binding site (Tsien et al., 1987). However, MD studies on calcium channels have been hindered by the lack of inaccurate calcium models in classical MD.TRPV channels are excellent candidates for investigating calcium channel selectivity.The TRPV family has six members, and their structures share a similar tetramer scaffold, but have distinct calcium selectivity (Venkatachalam and Montell, 2007).Among these, only TRPV5 and TRPV6 are thought to be Ca^2+^-selective (P_Ca_:P_Na_ > 100). Other members of this family, including the capsaicin receptor and thermosensor TRPV1, are considered nonselective cation channels (P_Ca_:P_Na_ < 10) (Owsianik et al., 2006). Furthermore, the open-state structures of Ca^2+^-selective TRPV6 and non-selective TRPV1 are available (Cao et al., 2013; McGoldrick et al., 2018), providing great opportunities to investigate their ion binding, permeation, and selectivity using MD simulations. In closed-state crystal rTRPV6 structures, three Ca^2+^ binding sites in the selectivity filter (SF) were identified, formed by D541, T539, and M570, respectively (Saotome et al., 2016). In the open-state cryoEM hTRPV6 structure, only one Ca^2+^-binding site near D542 in the middle of the SF was identified (McGoldrick et al., 2018). The constriction site in the SF of TRPV1 is formed by a simple glycine and has a rather spacious vestibule above it. Compared with non-selective TRPV1, Ca^2+^-selective TRPV6 has a longer and narrower SF formed by four residues, ^539^TIID^542^. How the differences in SF structures between TRPV6 and TRPV1 determine their distinct Ca^2+^ selectivity remains to be elucidated.

In this study, we conducted MD simulations for TRPV1 and TRPV6 using a recently developed multi-site Ca^2+^ model (Zhang et al., 2020), which is more accurate in describing interactions between Ca^2+^ and proteins and has been successfully used to simulate multiple Ca^2+^ channels (Liu et al., 2021; Schackert et al., 2022; Zhuang et al., 2022), to reveal the ion binding and permeation mechanisms through TRPV channels.Our results showed that despite the similar overall protein scaffolds, the differences in the SF structure resulted in distinct ion binding and permeation patterns in TRPV6 and TRPV1. The binding competition and number of binding sites in SF could determine Ca^2+^ selectivity under physiological conditions.

## Results

### The flexibility of the pore helix influences the cation permeation of TRPV channels

In our atomistic simulations of the TRPV systems (Fig. 1A), we applied position restraints to maintain the open conformation of the TRPV channels and found that various position restraints led to distinct permeabilities, particularly those restraints on the SF (Fig. 1B). In a 500-ns simulation of the pore domain of TRPV6, when the entire pore domain was restrained to the open-state cryoEM structure (termed “Restrained Protein”), the narrow and rigid SF, with the entrance region formed by D542 having a radius of less than 1 Å (Fig. 1C), was not permeable to cations and no single Na^+^ permeation was observed (Fig. 1D). Removal of the restraints on the SF loop (termed “Flexible SF”) gave limited flexibility to the SF, and the radius was not significantly dilated (Fig. 1C). No Na^+^ permeation was observed in a 500-ns simulation under such conditions for TRPV6 (Fig. 1D). To further relax the SF region, the loops and pore helix between S5 and S6 were made fully flexible (termed “Flexible Pore Helix”). Under such a condition, the continuous permeation of Na^+^ was observed (Fig. 1D). The radius of the pore was dilated to approximately 1.5 Å at the SF, and the fluctuation increased compared with that in the more restrained situations (Fig. 1C), allowing the passage of sodium through the SF (Fig. 1D). The conductance calculated from the simulation with the flexible pore helix (~70 pS) is comparable to the experimental value of 42–58 pS (Yue et al., 2001). These results highlight the importance of SF flexibility for ion permeation.

**Figure 1.**
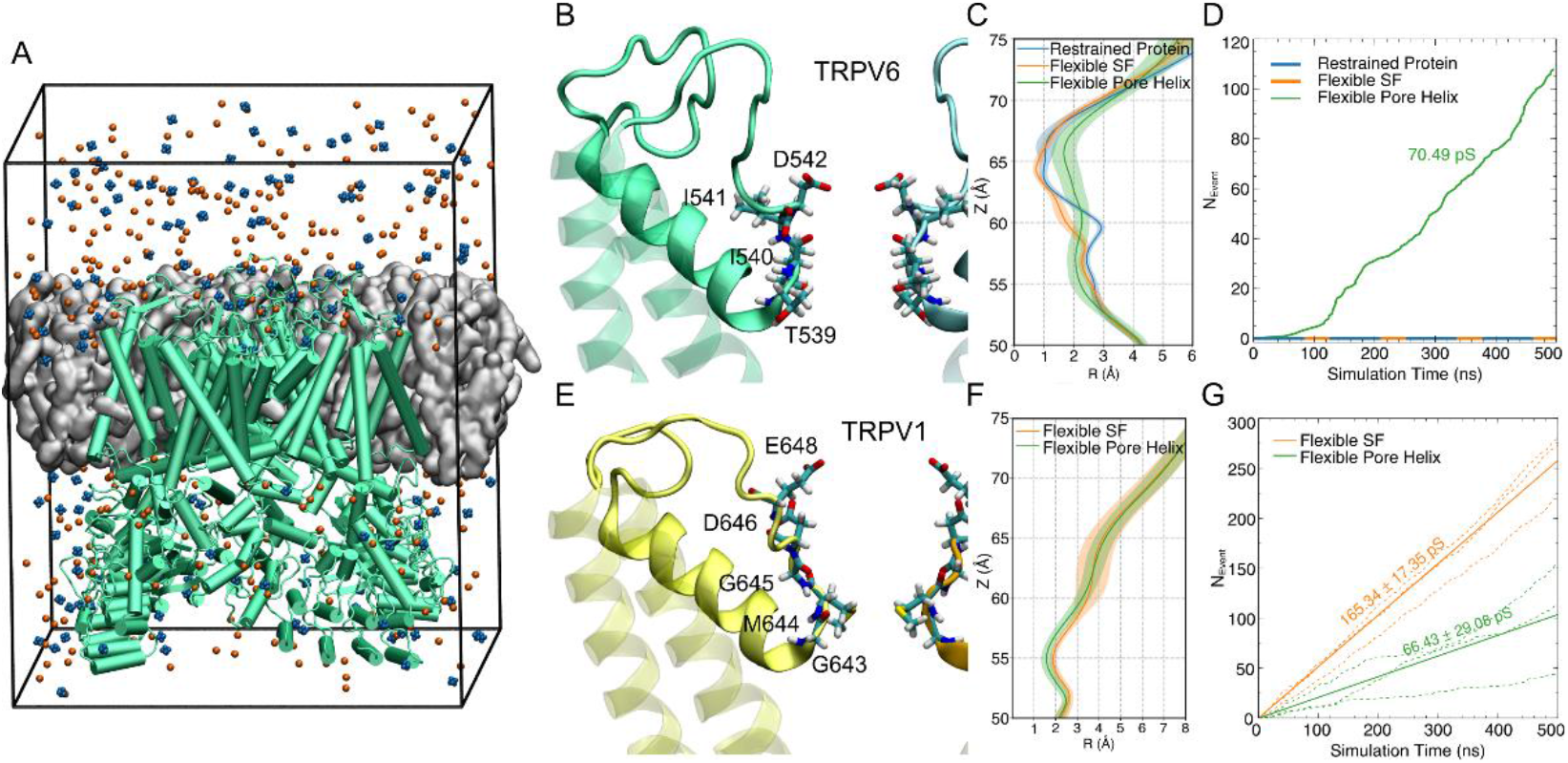
The flexibility of the pore helix influences the cation permeation of TRPV channels. **(A)** A representative simulation system, which contains the full TRPV6 (green cartoon) embedded in a POPC bilayer (gray surface). The multi-site Ca^2+^ ions are shown as blue spheres and Cl^-^ as orange spheres. Water molecules are filled in the box but not shown for clarity. **(B)** The structure of the SF region of TRPV6. The backbone of protein is shown as a cartoon with the pore helix and loops shown as solid and other parts as transparent. Key residues forming the SF are shown as licorice and labeled. Only two chains are shown for clarity. **(C)** The average pore radius of the SF region of TRPV6 during simulations with different restraints. The standard deviation is shown as the shaded area. **(D)** The cumulative number of Na^+^ permeation events (*N*_*Event*_) during simulations with different restraints. The ion conductances calculated based on the number of permeation events were labeled. **(E) to (G)** are similar to (B) to (D), respectively, but for TRPV1. The solid lines correspond to the average conductance from three independent simulation trajectories represented by dash lines.

In TRPV1, the flexibility of the pore loops and helix (Fig. 1E) also influenced the radius at SF during the simulations, thus impacting cation permeation. The open-state structure used in this study was solved using the spider toxin peptide DkTx and the vanilloid agonist RTX, whose binding to TRPV1 induces pain in humans. DkTx binds to the extracellular part of the channel and induces a conformational change in the loop between S5 and S6 (Gao et al., 2016). Contrary to TRPV6, the less restrained SF of TRPV1 showed a slightly smaller radius (Fig. 1F), whereas the number of permeation events dropped by more than half (Fig. 1G), resulting in a conductance close to the experimental value (62.8 pS) (Samways and Egan, 2011).

Simulations of TRPV6 and TRPV1 with different restraints applied to the pore loops and pore helix revealed that the flexibility of this region is essential for cation permeation. Based on the above benchmarking, position constraints were only applied to the transmembrane helices in the following production simulations, leaving the loop region and pore helix between the transmembrane helices fully flexible.

### Cytosolic domains of TRPV channels regulate ion permeation as well

To evaluate the role of the cytosolic domain of TRPV channels in ion permeation, simulations with and without cytosolic domains at 150 mM Ca^2+^ or 150 mM Na^+^ ion concentrations were conducted. As TRPV1 and TRPV6 have similar cytosolic domains with six ankyrin repeats and β-sheet structures, we chose TRPV6 as the simulation object because the cytosolic domain was more complete in the solved TRPV6 structure. In both simulations with Ca^2+^ or Na^+^, the presence of the cytosolic domain resulted in a decline in ion conductance and an increase in the permeation time of ions through the SF and gate regions (Fig. 2). Comparing the median permeation time, the increase was similar in the SF and gate regions, but was greater for Ca^2+^ than for Na^+^ (1.62-fold increase in SF, 1.42-fold increase in gate for Na^+^, 2.21-fold increase in SF, and 2.52-fold increase in gate for Ca^2+^). Ion density maps showed that Na^+^ mainly binds to the transmembrane domain and there is no strong binding site in the cytosolic domain (Fig. 2C). Thus, the cytosolic domain is speculated to influence Na^+^ permeation mainly by steric hindrance. In contrast, Ca^2+^ binds specifically to both the transmembrane and cytosolic domains (Fig. 2F). Multiple binding sites were observed near the residues E591 and D635 in the cytosolic domain. These electrostatic sticky sites act together with steric hindrance to regulate the conductance of ions more significantly for Ca^2+^ than for Na^+^.

**Figure 2.**
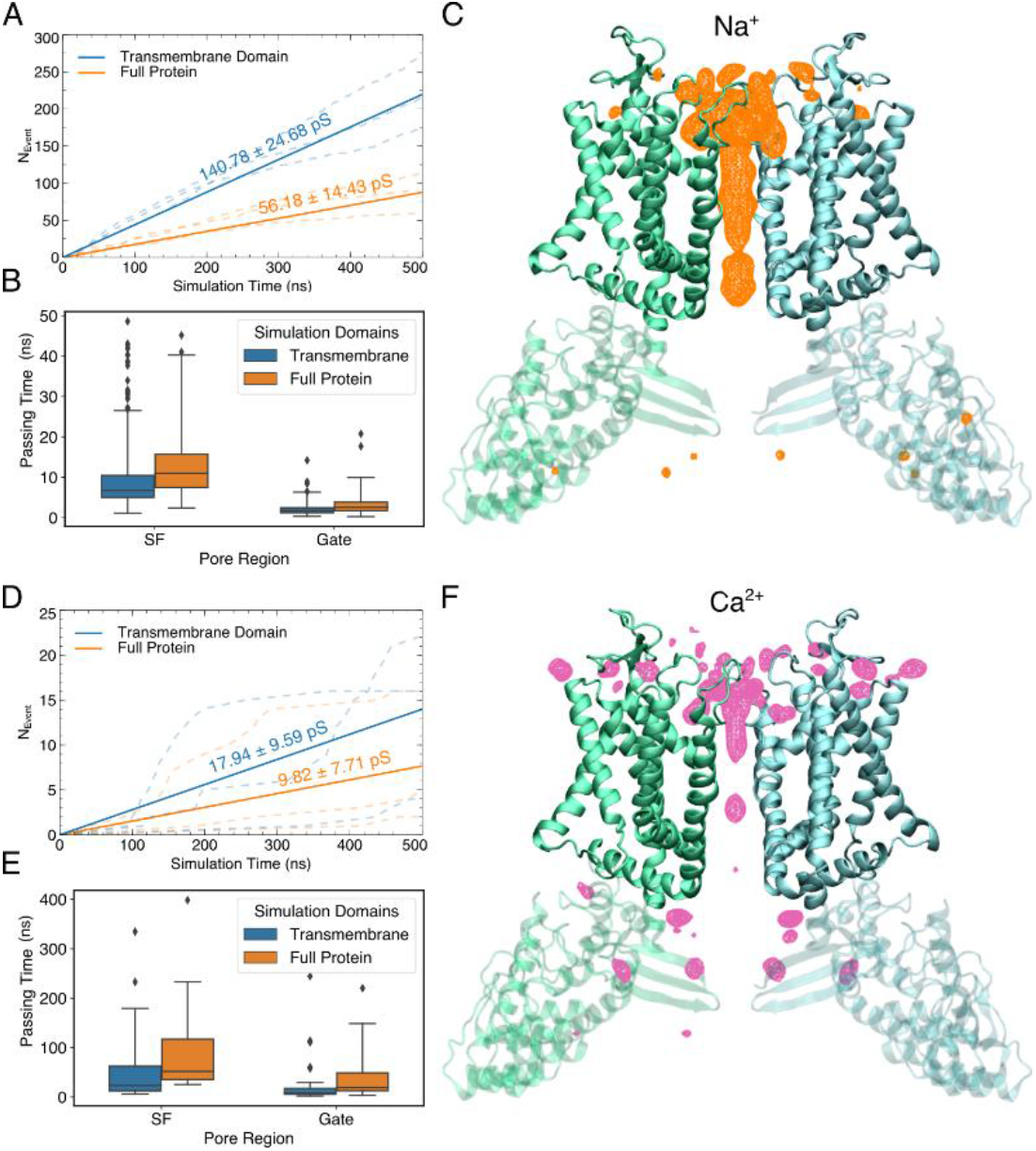
The cytosolic domain of TRPV6 regulates the permeation of cations. **(A)**The cumulative number of Na^+^ permeation events (*N*_*Event*_) in simulations with (Full Protein) and without cytosolic domain (Transmembrane Domain). The ion conductances calculated based on the number of permeation events were labeled. The solid lines correspond to the average conductance from three independent simulation trajectories represented by dash lines. **(B)** The time of Na^+^ passing through the SF and gate region during permeation in simulations with and without cytosolic domain. **(C)** The density map of Na^+^ calculated using trajectories of “Full Protein” simulations for TRPV6. The density is shown as a meshed isosurface. The backbone of the channel is shown as cartoon with the transmembrane domain shown as solid and the cytosolic domain as transparent. Only two chains are shown for clarity. **(D) to (F)** are similar to to (C), respectively, but for simulations with Ca^2+^ instead of Na^+^.

Steric hindrance and electrostatic sticky sites in the cytosolic domain are less likely to change the permeating pattern, including the permeating path, interactions between permeating ions and proteins, and interactions between permeating ions in the transmembrane domain. Considering the questions of interest and computational costs, we chose to include only the transmembrane domain of the TRPV channels in the following simulation systems for the ion binding and permeation study through the transmembrane pore.

### Differences in Ca^2+^-binding sites and knock-off permeation of TRPV1 and TRPV6

Our μs-scale simulations showed different binding sites for Ca^2+^ in the SF regions of TRPV6 and TRPV1. In TRPV6, two types of Ca^2+^-binding sites were identified: site S at the center of the SF and the peripheral binding site P (Fig. 3A & B). Binding site S can be subdivided into two sites: S1 and S2. S1 is located at the entrance of the SF and Ca^2+^ interacts with the D542 side chain. S2 is located in the lower SF, and Ca^2+^ mainly interacts with T539. Ca^2+^ at the binding site P mainly interacts with the negatively charged residue E535. Covering all binding sites, we defined the SF region of TRPV6 as the region formed by residue T539 to D542. The average number of Ca^2+^ ions in the SF region is 4.39. Notably, the binding sites observed in our simulations were not identical to the ion densities identified in previous structural studies. Compared with the closed-state crystal rTRPV6 structure (Saotome et al., 2016), our simulations showed two binding sites at the SF with higher electron density in the crystal structure, but the third binding site with lower electron density in the cavity was not obvious in our simulations. The open-state cryo-EM hTRPV6 structure solved the binding site located in the middle of the SF, but no Ca^2+^ density was observed at the entrance of the SF (McGoldrick et al., 2018). These differences between our simulations and existing structures can be partly explained by the gating state and flexibility of SF during ion permeation. The crystal structure was in a closed state, whereas it was difficult to capture the less stable ion binding sites in the cryoEM structure; therefore, our simulations provide more details on the ion distribution while permeating. In addition, the transmembrane potential applied in our simulations may cause a shift in the ion-binding site compared to the experimental observation under transmembrane potential-free conditions.

**Figure 3.**
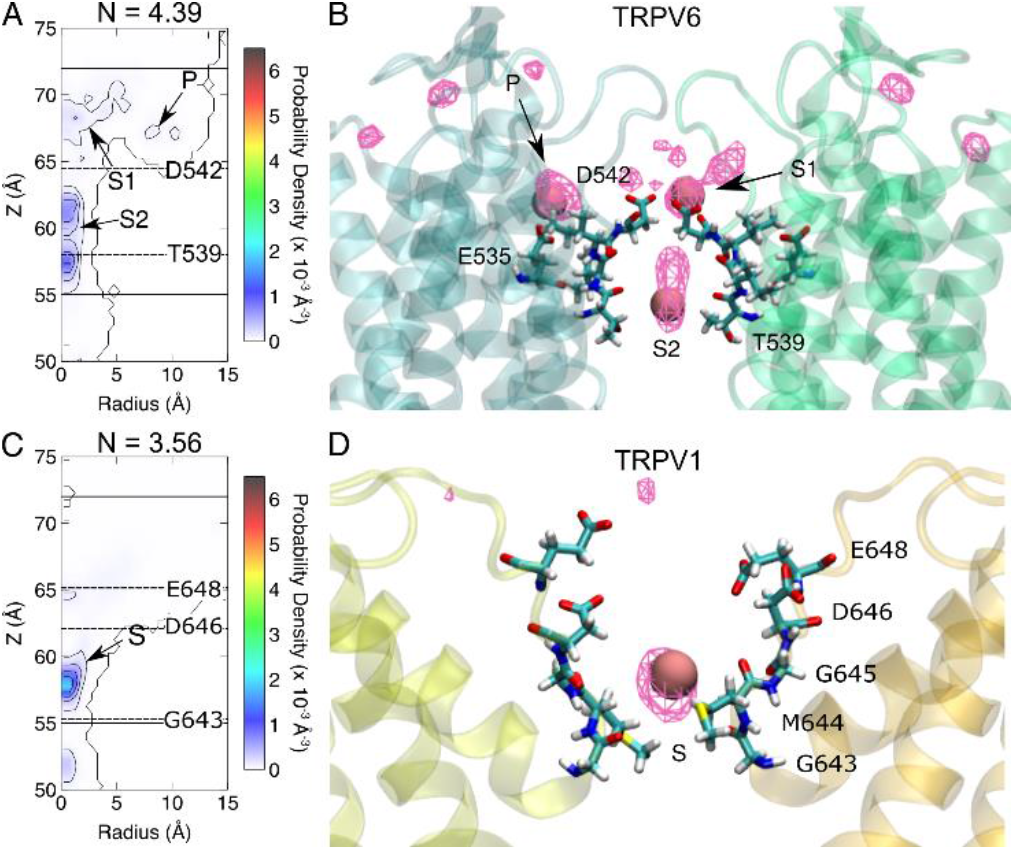
The difference in Ca^2+^ binding at the SF of TRPV channels leads to different ion permeation patterns. **(A)** The contour plot of Ca^2+^ density on the R-z plane of the SF region of TRPV6. Binding sites are indicated by arrows and labeled. The Cα positions of key residues are indicated by dashed lines. The boundaries of the SF are indicated with solid lines. N is the average number of Ca^2+^ in the SF region. The representative snapshot of Ca^2+^ binding at the SF region. The average ion density map is shown as a meshed isosurface. Binding sites are indicated by arrows and labeled. Residues forming the binding sites are shown as licorice. The Ca^2+^ is shown as a pink sphere. The backbone of protein is shown as a transparent cartoon. Only two chains are shown for clarity. **(C)** and **(D)** are similar to (A) and (B), respectively, but for TRPV1.

In TRPV1, only one stable Ca^2+^ binding site at SF was observed, which was located above the narrowest site formed by G643 (Fig. 3C and D). Ca^2+^ bound to this site mainly interacts with G643, M644, and G645. No peripheral Ca^2+^-binding site corresponding to site P in TRPV6 was observed in TRPV1. To be comparable to TRPV6, we defined the TRPV1 SF region in a similar way to that of TRPV6, including the wide extracellular vestibule and the narrowest site at G643. The average number of Ca^2+^ ions in this SF region was approximately 3.56, which is less than that in TRPV6.

Different Ca^2+^ binding sites in the SF region lead to different modes of permeation of TRPV6 and TRPV1. In both TRPV channels, a Ca^2+^ transport process consistent with a previously proposed knock-off mechanism was observed (Tsien et al., 1987).Continuous permeation of Ca^2+^ requires the entry of Ca^2+^ into the SF region to knock off Ca^2+^ at the binding site. However, in TRPV6, since two Ca^2+^ ions are bound at the SF, permeation is characterized by sequential knocking and concerted transport of both Ca^2+^ in the SF. As shown in Fig. 4A, when Ca^2+^ approaches the binding site S1 from the extracellular site, it knocks the Ca^2+^ bound at S1 by electrostatic repulsion. Sequentially, the Ca^2+^ bound at S1 knocks the nearby Ca^2+^ bound at S2, leading to the dissociation of Ca^2+^ to enter the channel cavity for permeation. Meanwhile, the Ca^2+^ bound at S1 moves to S2, and then S1 is occupied by a newly entering Ca^2+^ from the extracellular site. The three Ca^2+^ ions move in concerted and synergic ways to permeate. In contrast, for TRPV1, the calcium ion permeation process involves only one bound Ca^2+^ and one Ca^2+^ knocked out from the extracellular side, with no sequential knocking or concerted movement between multiple ions in the SF (Fig. 4B). However, it should be noted that here we only focus on the ions in the SF, while the ions in the cavity would probably be involved in the concerted permeation as well, as observed in the work by Ives et al. (Ives et al., 2022). Therefore, multiple ion binding and concerted permeation appears to be a universal phenomenon in the TRPV channels.

**Figure 4.**
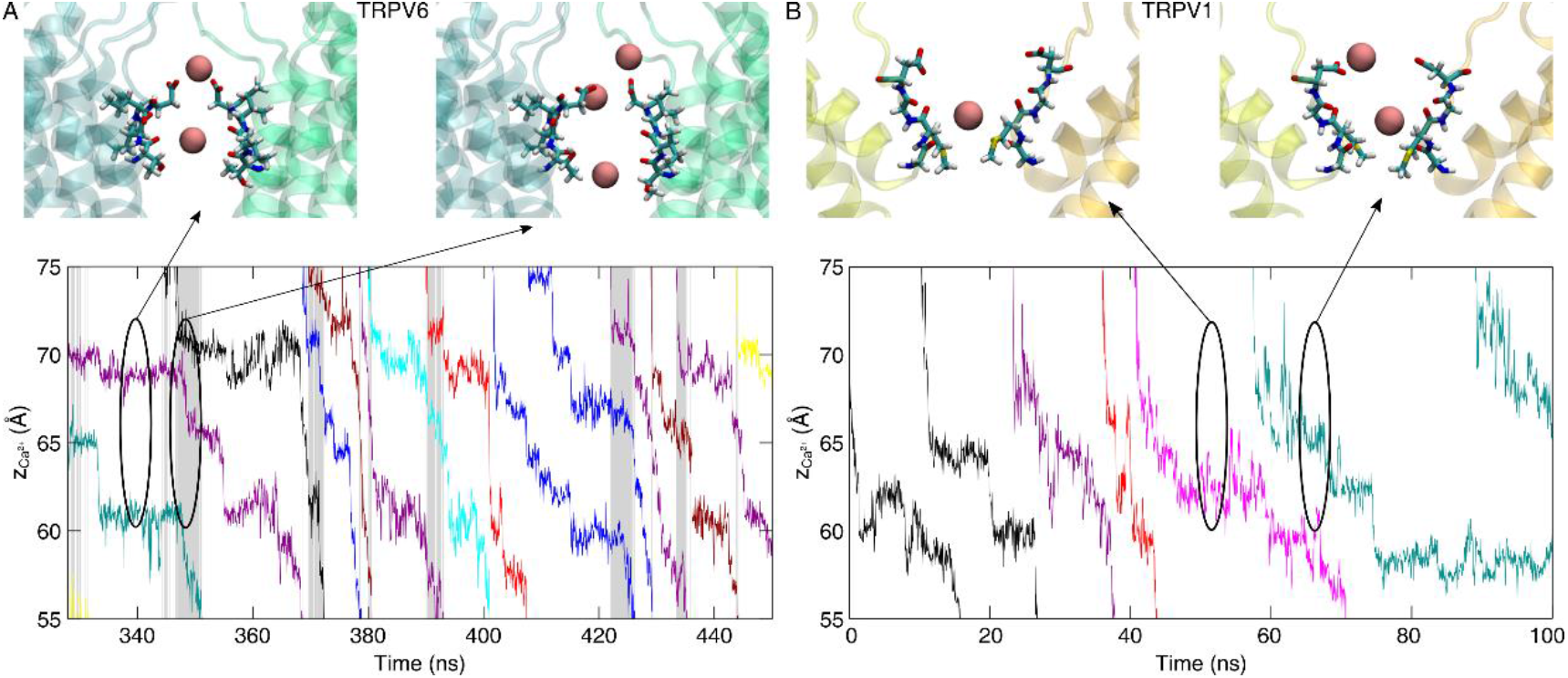
Different modes of Ca^2+^ permeation in TRPV6 and TRPV1. **(A)** The Evolution of the z-coordinates of permeating Ca^2+^ in TRPV6. Different permeating Ca^2+^ are shown in different colors. Shaded areas indicate the simultaneous 3-Ca^2+^-occupy in the SF region. The representative snapshot of Ca^2+^ permeating the SF region. Residues forming the SF are shown as licorice. The Ca^2+^ is shown as a pink sphere. The backbone of protein is shown as a transparent cartoon. Only two chains are shown for clarity. **(B)** similar to (A), but for TRPV1.

### The dehydration of Ca^2+^ is slightly different in TRPV6 SF than in TRPV1

The dehydration of permeating Ca^2+^ in the SF region has similarities, but also shows discrepancies between TRPV6 and TRPV1. Ca^2+^ is slightly dehydrated in the SF region of both TRPV6 and TRPV1, but dehydration in TRPV6 occurs in a longer region and is more significant than that observed in TRPV1. In TRPV6, at most 1.5 water molecules in the first solvation shell were dehydrated, and this dehydration was mainly replaced by oxygen atoms on the side chains of charged residues (D542 and E535) in the SF region (Fig. 5A). In TRPV1, dehydration of permeating Ca^2+^ at the SF in the TRPV1 channel occurred only at the narrowest point formed by G643, where Ca^2+^ was coordinated by the oxygen atoms of the protein backbone. At most, one water molecule in the first solvation shell of Ca^2+^ was dehydrated in the SF of TRPV1, which was slightly weaker than that in TRPV6 (Fig. 5B). It should be noted that although partial dehydration of Ca^2+^ occurs at the negatively charged residues E648 and D646 above the SF of TRPV1, the pore radius at this location is larger than the radius of the first solvation shell formed by coordinated waters around Ca^2+^ (~ 4 Å). Therefore, we consider the dehydration of the calcium ion here not due to spatial constraints but due to the stronger electrostatic attraction between the Ca^2+^ and the negatively charged residues. In addition, the calcium ions dehydrated here do not necessarily participate in the permeation, so ion dehydration here was not considered as dehydration in permeation in this study. Therefore, the dehydration of permeant Ca^2+^ mainly occurs around the constriction site in the upper SF of TRPV6, whereas in TRPV1, this mainly occurs around the constriction site in the lower SF. When dehydration occurs at these constriction sites, there are always oxygen atoms of proteins that can replace water oxygens to coordinate with the permeant Ca^2+^ (Fig. 5).

**Figure 5.**
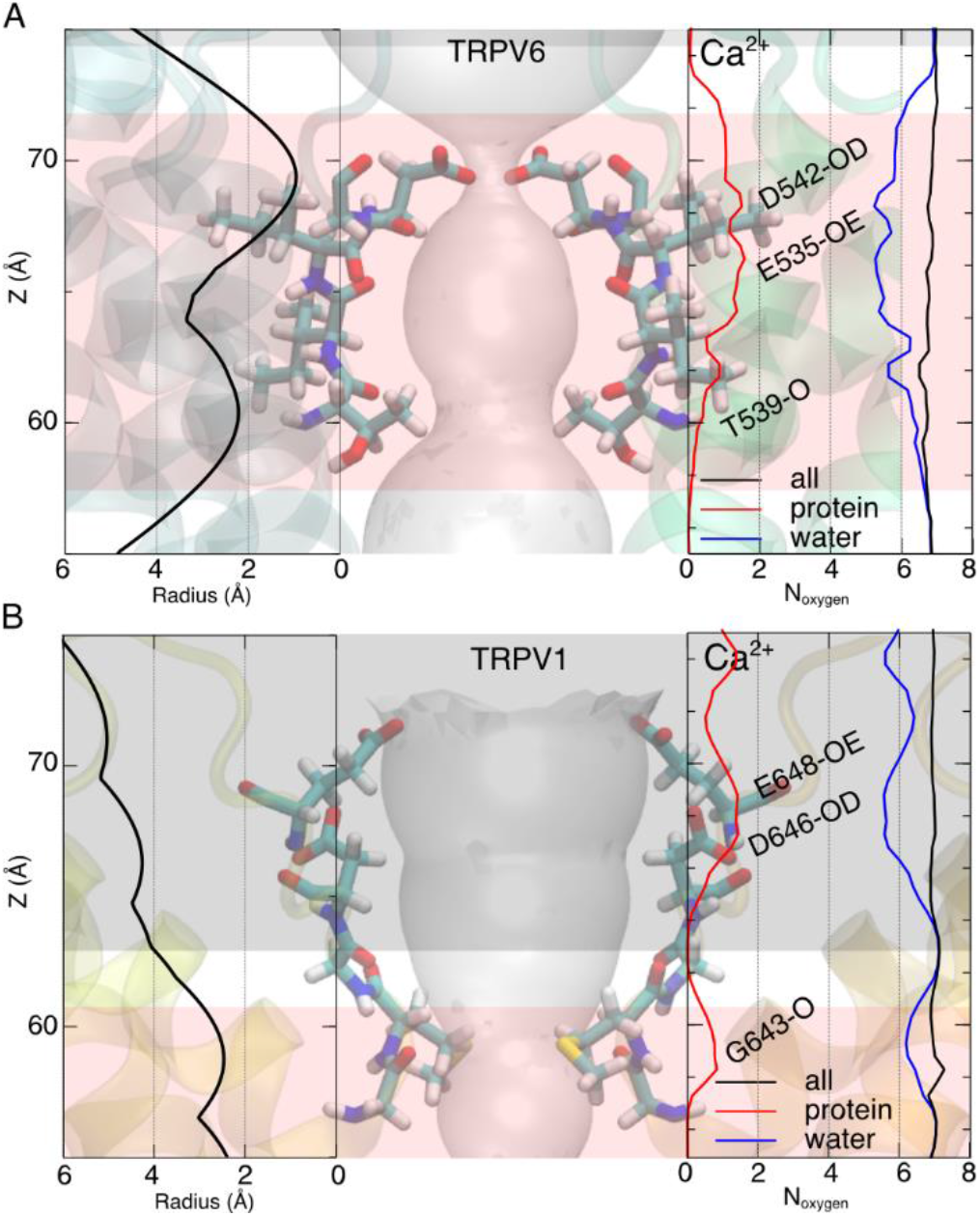
The dehydration of calcium is more significant in TRPV6 SF than in TRPV1. **(A)** The dehydration of Ca^2+^ during the permeation of TRPV6. The left panel shows the pore radius along the pore axis of the open-state TRPV6. The right panel shows the total number (black) of coordinated oxygen atoms around the Ca^2+^ within the pore, and the contributions from protein (red) and water (blue) respectively. The significant contributions from protein oxygen are marked with corresponding protein residue IDs containing the oxygen. The transparent background shows the pore profile within the open-state TRPV6 structure. The pore region where the radius is larger than 4 Å is marked with dark shadows. The SF region is highlighted by the red shadow. **(B)** similar to (A), but for TRPV1.

### The blockage effect of Ca^2+^ on monovalent current and valence selectivity

In simulations with di-cationic solutions, the blockage effect of divalent ions on monovalent currents was observed for both TRPV6 and TRPV1. The conductance of the channel decreased significantly when Ca^2+^ was added to the Na^+^ solution (Table 1). The stronger blockage effect observed in simulations with TRPV6 than with TRPV1 (19.13-fold decrease in TRPV6 vs. 2.47-fold decrease in TRPV1) is consistent with experimental observations (Vennekens et al., 2000; Samways and Egan, 2011), which may be related to the stronger Ca^2+^ selectivity of TRPV6 over TRPV1 (Gillespie and Boda, 2008) (Table 1). Despite the blockage effect of Ca^2+^, permeation events of Na^+^ were still observed in our simulations. The Na^+^ permeation patterns differed between TRPV6 and TRPV1. In TRPV6, Na^+^ is not permeable when Ca^2+^ binds to the SF. Only at the time interval between Ca^2+^ permeation when one Ca^2+^ leaves the binding site S1 can the nearby Na^+^ take the opportunity to bind to this site and cut in line to permeate (Fig. 6A and B). Since the binding Ca^2+^ moves in concert during permeation, another Ca^2+^ will occupy the binding site soon after one Ca^2+^ leaves, so the time interval available for Na^+^ to cut in line is rather limited. Thus, in TRPV6, calcium ions show a very strong blocking effect on monovalent currents. In contrast, the vestibule above the SF region in TRPV1 is rather spacious; when one Ca^2+^ is present in this region, Na^+^ can bypass it and permeate (Fig. 6C and D). This bypass permeation pattern of Na^+^ in TRPV1 is more likely to occur than the cut-in-line pattern in TRPV6. Therefore, the blockage effect of Ca^2+^ on monovalent currents is weaker in TRPV1 than in TRPV6.

**Table 1.** The overall conductance in simulations with different concentrations of cations.

| Ion<br>Channel | Calcium<br>(150 mM) | Sodium<br>(150 mM) | Sodium + Calcium<br>(150 mM + 150 mM) |  |
| --- | --- | --- | --- | --- |
| | Conductance (pS) | | Conductance (pS) | $N_{Event}^{Ca^{2+}}:N_{Event}^{Na^{+}}$ |
| TRPV6 | $17.94 \pm 9.59$ | $140.78 \pm 24.68$<br>(42–58*) | $7.36 \pm 4.59$ | 2.06 |
| TRPV1 | $11.96 \pm 7.99$<br>(15.2**) | $66.43 \pm 29.08$<br>(62.8**) | $26.92 \pm 16.32$ | 7.38 |

\*(Yue et al., 2001)
\*\*(Samways and Egan, 2011)
The experimental conductance is shown in the brackets.

**Figure 6.**
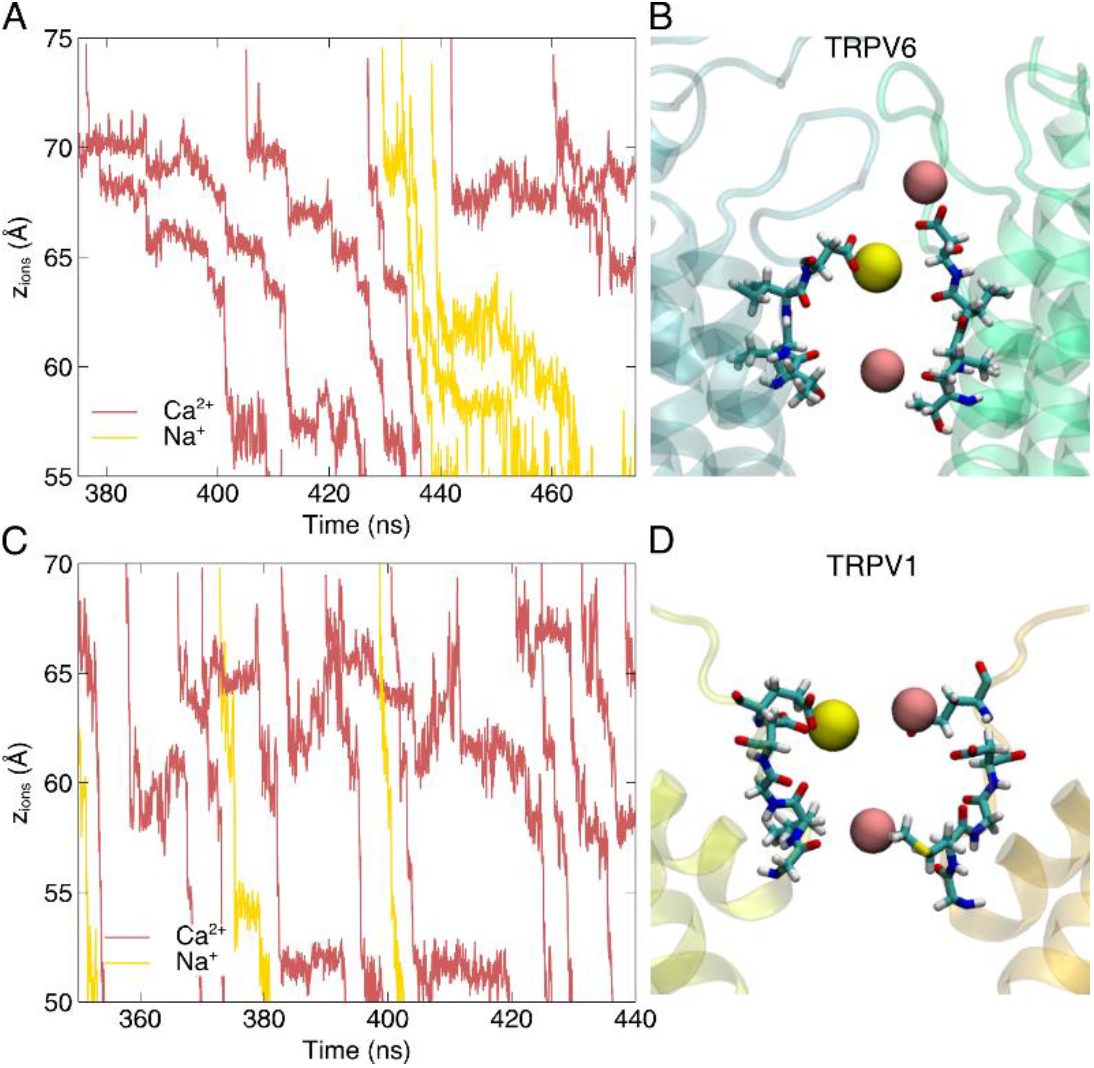
Na^+^ shows different permeation behavior in TRPV6 and TRPV1 in the presence of Ca^2+^. **(A)** Evolution of the z-coordinates of TRPV6-permeating cations in the di-cationic simulations. **(B)** Representative conformation during Na^+^ permeation of TRPV6. The backbone of protein is shown as a transparent cartoon. Residues forming the SF region are shown as licorice. Only two chains are shown for clarity. Ca^2+^ is shown as the pink sphere and Na^+^ as the yellow sphere. **(C)** similar to (A), **(D)** similar to (B), but for TRPV1.

In the di-cationic systems, TRPV6 and TRPV1 showed slight valence selectivity in our MD simulations. As shown in Table 1, TRPV1 showed a slight preference for Ca^2+^ permeation over Na^+^ (~7:1), which is consistent with the previous conclusion that TRPV1 is not Ca^2+^ selective. However, for TRPV6, we did not observe any significant Ca^2+^ selectivity either. The ratio of permeation events was approximately 2:1 (N_Ca2+_:N_Na+_), which is inconsistent with the prediction that the permeability ratio P_Ca_:P_Na_ is greater than 100:1. This is discussed further in the next section.

### Binding site competition at SF entrance facilitates Ca^2+^ selectivity/Na^+^ blockage in TRPV6

In the vestibule above the SF, Ca^2+^ and Na^+^ compete to occupy binding site S1. In our di-cationic simulations, a significant decrease in the density of Na^+^ in the SF region of both TRPV6 and TRPV1 was observed compared to the system with only Na^+^ as a cation, indicating that Ca^2+^ repels Na^+^ from this region (Fig. 7A and C). In TRPV1, Na^+^ density in the entire SF region decreased (Fig. 7D). In contrast, in TRPV6, Na^+^ density showed a slight increase at the peripheral binding site P (red circle in Fig. 7B). The differentiation of Ca^2+^ and Na^+^ binding in TRPV6 SF was observed by a detailed inspection of their 3D densities. Ca^2+^ mainly binds to the central binding site S, including S1 and S2, whereas Na^+^ is preferentially bound at the peripheral binding site P (Fig. 7E). Ion transition probability analysis between these binding sites showed that for both Ca^2+^ and Na^+^, the ions bound at S2, which is the indispensable site during permeation, were all transferred from the S1 binding site in the single cationic systems, indicating that S1→S2 is the major permeation path. In the di-cationic systems, all of the Ca^2+^ ions still followed this single permeation path, while for Na^+^, 81% of the permeation events followed the S1→S2 path and 19% followed a new P→S2 path. Most of the permeating ions once bound at site P moved to the S1 binding site for further permeation to S2, but some of them still moved directly from the binding site P to S2 (Fig. 7F). This inequality in the transition rate to S2 indicates that the ions bound at site S1 have a significantly higher permeation probability, and those bound at site P are less involved in direct permeation. In the TRPV6 di-cationic simulation system, Ca^2+^ dominantly occupied the central binding sites, having a higher probability of permeation, while Na^+^ was expelled to the peripheral binding sites P, having a lower probability of permeation. This binding site competition and differentiation between cations of various valences may also contribute to selectivity for Ca^2+^ or blockage of Na^+^ in TRPV6.

**Figure 7.**
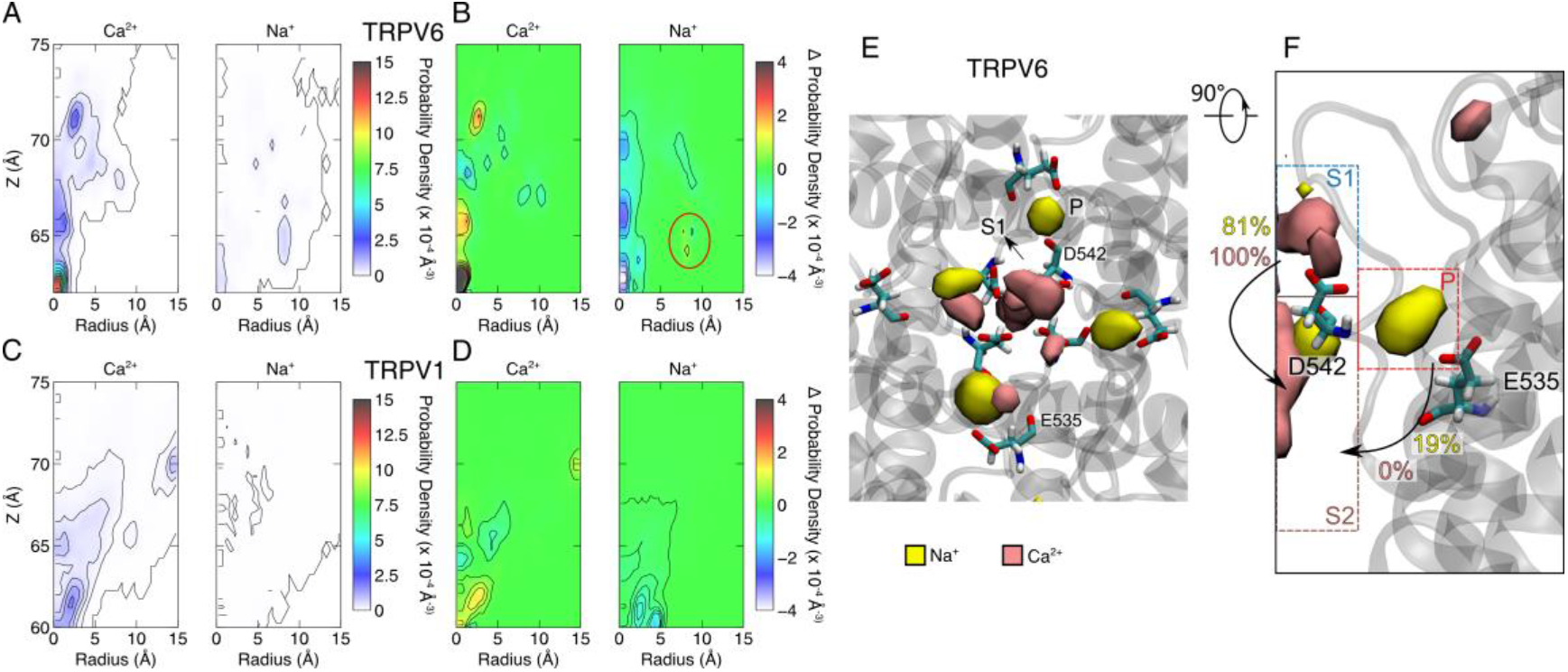
The binding site differentiation between Ca^2+^ and Na^+^ in the SF region of TRPV6. **(A)** The contour plot of cation density on the R-z plane of the SF region of TRPV6. **(B)** The difference in cation density calculated by subtracting the density of the one-cation-species system from the density of the di-cationic system. The peripheral (P) binding site is indicated by the red circle. **(C)** similar to (A), **(D)** similar to (B), but for TRPV1. **(E)** Top view of the density map of cations in the SF region of TRPV6. The backbone of the protein is shown as the gray transparent cartoon. The density map is shown as the solid isosurface. The binding sites and key residues are labeled. Only two chains are shown for clarity. **(F)** Side view of (D). Only one chain is shown. Binding sites are indicated by dashed rectangles. The transition probabilities of S1→S2 and P→S2 are labeled on the corresponding arrows, with yellow for Na^+^ and pink for Ca^2+^.

## Discussion

Various Ca^2+^-permeable channels showed significantly different Ca^2+^ selectivity, with reported permeability ratios over monovalent cations ranging from ~1:1 to ~1000:1, which are required for these channels to execute their respective biological functions (Sather and McCleskey, 2003). How ion selectivity of various degrees is achieved has long been an intriguing question in the field of ion channel research, which requires detailed structural and dynamic insights at the atomic level. Homologous ion channels that show distinct Ca^2+^ selectivity and whose structures have been resolved, such as the TRPV channels studied in this work, provide an excellent opportunity to address this issue.

Our MD simulations of cation permeation in Ca^2+^-selective TRPV6 and non-selective TRPV1 showed that cationic binding competition in multiple Ca^2+^ binding sites in the SF is probably the main cause of Ca^2+^ selectivity. From the permeation trajectory analysis, it was observed that Ca^2+^ showed a much stronger binding preference than Na^+^ at the SF entrance of both TRPV6 and TRPV1. In the presence of Ca^2+^, Na^+^ ions were expelled from the central binding site (S1) at the SF entrance (Fig. 7). Although Na^+^ can still occupy the lateral binding site P near the SF entrance of TRPV6, the binding site P does not lead to efficient permeation into the SF. The preferential binding of Ca^2+^ at the SF entrance can generate a blocking effect on Na^+^, which can enhance the permeation probability of Ca^2+^ over Na^+^ for both TRPV6 and TRPV1.

The number of binding sites in SF can generate distinct permeation patterns. With one Ca^2+^ binding site in the SF, Na^+^ can still frequently find their chance to enter the SF, as observed in TRPV1. A possible factor that can enhance Ca^2+^ selectivity is the presence of multiple Ca^2+^-binding sites in the SF, as observed in the SF of TRPV6. When the lower Ca^2+^ binding site (S2) is occupied by Ca^2+^, the upper binding site (S1) can be occupied by either Ca^2+^ or Na^+^, with a stronger binding preference for Ca^2+^.Meanwhile, only when S1 is occupied by Ca^2+^ does the Ca^2+^ at the S2 site in the SF exert a stronger electrostatic driving force to overcome the permeation energy barrier under physiological conditions. Therefore, multiple Ca^2+^ ions must line up and march in a concerted manner to permeate more efficiently. Consequently, multiple Ca^2+^ binding sites lead to a concerted permeation manner, which is a characteristic that discriminates the Ca^2+^ permeation in TRPV6 from TRPV1, as also observed by Ives et al. (Ives et al., 2022). In fact, increasing the number of Ca^2+^-binding sites in channel pores has long been proposed to improve channel selectivity for Ca^2+^ (Tsien et al., 1987). Under physiological conditions, this concerted permeation method only allows Na^+^ permeation when both Ca^2+^ in the SF are away, which is much rarer than the case with only one Ca^2+^ binding site. Therefore, the multiple Ca^2+^ binding sites within SF can reject Na^+^ permeation more efficiently than in TRPV1, resulting in higher Ca^2+^ selectivity. However, in our simulations, although we observed concerted Ca^2+^ permeation, Na^+^ permeation events still occurred in TRPV6. We attribute this to the fact that a high transmembrane voltage (500 mV) was applied in our MD simulations to obtain sufficient statistics, which would generate a much stronger driving force for cations to permeate than under physiological conditions. Thus, monovalent cations can probably still push Ca^2+^ through SF, resulting in a lower valence selectivity than expected. Indeed, Ives et al.’s work observed similar behavior for multiple TRPV channels, as well as that the selectivity increases in TRPV5 with decreasing the transmembrane potential (Ives et al., 2022).

Ca^2+^ permeates through the SF of both TRPV6 and TRPV1 in a slightly dehydrated manner, and dehydration occurs in a more extended region of TRPV6, suggesting that the permeation pathway is more constricted for hydrated Ca^2+^ in the SF of TRPV6 than in TRPV1. Fortunately, the dehydration of Ca^2+^ in TRPV6 occurs around the charged residues in the SF, and the oxygen atoms of the charged residues can take the role of coordination with the permeant Ca^2+^, thus lowering the dehydration free energy barrier. In contrast, for TRPV1, the steric constriction site is below the electrostatic attraction site in SF. Although TRPV6 and TRPV1 utilize electrostatic and steric interactions to attract and bind Ca^2+^, they have different strategies to regulate ion permeation in the SF. The distinct permeation mechanisms of TRPV6 and TRPV1 are based on their distinct SF structures. In the Ca^2+^-selective channel TRPV6, two adjacent binding sites line up in the narrow and long SF regions, consisting of negatively charged D542 and polar T539. This SF structure, which is approximately 2 nm in length, enables Ca^2+^ to bind partially dehydrated and allows two Ca^2+^ ions to bind simultaneously in the SF region. In contrast, the SF of TRPV1 is much shorter and has a wide vestibule of negative electrostatic potential above the SF, with the narrowest region consisting of only one non-charged G643. This structure only allows for one Ca^2+^ binding site, and Ca^2+^ in the vestibule is flexible and less efficient in blocking the permeation of Na^+^.

Although conventionally called “knock-off” (Hess and Tsien, 1984; Tang et al., 2014), the concerted Ca^2+^ permeation through multiple binding sites in TRPV6 is similar to the knock-on permeation behavior in the selective Na^+^ and K^+^ channels, which possess two Na^+^ and four K^+^ binding sites in the SF region, respectively (Corry and Thomas, 2012; Köpfer et al., 2014). However, there are evident differences in the selectivity mechanisms of these channels. In the highly selective Ca^2+^ channels, such as TRPV6, slightly dehydrated Ca^2+^ ions occupy two binding sites in the SF that are farther away from each other (with a separation of 11 Å), and the determinative factor for ion selectivity is likely the difference in electrostatic binding affinity and driving force generated by different valences. In the highly selective K^+^ channels, fully dehydrated K^+^ ions sit next to each other (with a separation of about 3.4 Å), and the determinant factor for ion selectivity is the dehydration energy difference of various ions when entering the SF. The highly selective Na^+^ channels also showed two loosely packed Na^+^ binding sites in the SF (with a separation of 10 Å), where the determinative factor was the partial dehydration energy when Na^+^ entered the SF. In fact, the difference in the (partial) dehydration energy of K^+^ and Na^+^ entering the SF can also be viewed as the difference in the binding affinities of these monovalent ions to the SF.

Therefore, by studying two representative TRPV channels with distinct Ca^2+^ binding and permeation patterns, as well as relating the results to the existing knowledge of highly selective Na^+^ and K^+^ channels, we conclude that ion-binding competition at multiple binding sites in the SF of ion channels is required to generate high blocking effect or ion selectivity. Ion competition at one binding site can generate a moderate blocking effect and ion selectivity. To achieve high blocking or selectivity, multiple ion-binding sites with the same ion preference in SF are required. In such a scenario, the permeating ions would have to follow a concerted knock-on permeation behavior, which can simultaneously facilitate optimal ion permeability and selectivity. However, the selectivity may be transmembrane potential-dependent; a high voltage would abolish valence selectivity. In addition, our simulations showed that not only the specific state of the channel structure, but also the flexibility of the SF is essential for regulating ion permeability, as is the cytoplasmic domain. Therefore, to more quantitatively study ion permeation and valence selectivity under physiological conditions, complete ion channel structures in a realistic simulation setup, more accurate ion models that explicitly consider polarization, lower transmembrane voltage, and longer simulation time are desirable in future simulation studies.

## Methods

### Molecular dynamics simulations of TRPV channels

The open-state structure of hTRPV6 (PDB ID:6BO8) (McGoldrick et al., 2018) used in this study was solved using cryo-EM at a resolution of 3.6 Å. The missing loop between S2-S3 in this structure was modeled using Modeller (Šali and Blundell, 1993), utilizing the existing open structures (PDB ID:6BO8, 7K4A) (Bhardwaj et al., 2020) as the template. The whole protein structure and two truncated structures were used to build the protein-membrane simulation systems, retaining the channel pore domain (residue 475-588), and transmembrane domain (residue 317-608), respectively. For TRPV1, the DkTx/RTX-bound open-state structure solved using cryo-EM at a resolution of 2.95 Å was used (Gao et al., 2016). A truncated structure containing only the transmembrane domain (residue 423-713) was used for the simulations.

A POPC bilayer in the simulation system was built using the CHARMM-GUI Membrane Builder (Wu et al., 2014). Ions were added to the system to reach different target concentrations using the CHARMM sodium model and the multi-site calcium model developed by us recently (Zhang et al., 2020). The simulation systems were equilibrated using the standard CHARMM-GUI equilibration protocol before the production simulations (Jo et al., 2008). A transmembrane potential of 500 mV from the extracellular side to the cytosolic side was applied in all simulations. All simulations were performed using the CHARMM36m force field (Huang et al., 2017) and the TIP3P water model with a time step of 2 fs. The v-rescale thermostat (Hess et al., 2008) with a time constant of 0.5 ps and the Parrinello-Rahman pressure coupling (Parrinello and Rahman, 1981) with a time constant of 5 ps were used to maintain the temperature and pressure at 310 K and 1.0 bar during the simulations. The particle-mesh Ewald method (Darden et al., 1993) was used to calculate electrostatic interactions. The van der Waals interactions were smoothly switched off from 1.0 nm to 1.2 nm. Gromacs 2018.6 was used to run the MD simulations and trajectory analysis (Abraham et al., 2015).

Harmonic position restraints were applied to the α-carbon of the protein with a force constant of 1000 kJ mol^−1^ nm^−2^, while some regions in the SF were free without restraints in some simulations. In simulations with the pore domain of TRPV6, the pore loop-forming SF (residue 539-542) and pore helix regions (residue 514-552) were free. In simulations with the transmembrane domain of TRPV6, the loops between the transmembrane helices and pore helix regions were free (residue 350-378, 405-424, 447-449, 470-475, 514-552). In TRPV1 simulations, the loop formed SF (residue 643-656) and the loops between transmembrane helices and pore helices (residue 456-468, 501-510, 533-537, 557-561, 600-656) were free.

## Data analysis

The pore radii of the channels were calculated using HOLE (Smart et al., 1996). The channel conductance of permeating ions with charge Q_ion_ was calculated based on the number of permeation events (N_Event_) over the entire trajectory. A permeation event was defined as one ion permeating the membrane through a channel from the extracellular side to the cytosolic side. The conductance (C) of a trajectory of a certain time length (t) under transmembrane potential (V_tm_) was calculated as follows:

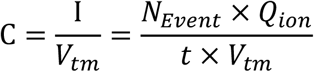

The probability density of the ions in the pores on the R-z plane was calculated as follows:

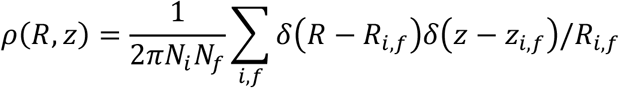

where *R* represents the distance from the central axis of the channel, *z* is the coordinate perpendicular to the membrane, *N*_*i*_ the number of ions in the system and *N*_*f*_ the number of frames recorded in the trajectory.

To calculate the permeation transition probability among the binding sites in the SF, we monitored the trajectories of all the ions that came from the SF and left binding site S2 to enter the cavity. The number of ions that followed the permeation path S1→S2 was designated as N_s1_, the number of ions that followed the permeation path P→S2 was N_P_, the transition probability of S1→S2 was calculated as 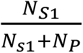, and the transition probability of P→S2 was 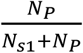.

